# Large genetic analysis of alcohol resistance and tolerance reveals an inverse correlation and suggests ‘true’ tolerance mutants

**DOI:** 10.1101/2023.10.09.561599

**Authors:** Maggie M. Chvilicek, Alexandra Seguin, Daniel R. Lathen, Iris Titos, Pearl N Cummins-Beebe, Miguel A. Pabon, Masa Miscevic, Emily A. Nickel, Collin B Merrill, Aylin R. Rodan, Adrian Rothenfluh

**Affiliations:** Department of Psychiatry, Huntsman Mental Health Institute, School of Medicine, University of Utah, Salt Lake City, USA; Neuroscience Graduate Program, University of Utah, Salt Lake City, USA; Molecular Medicine Program, School of Medicine, University of Utah, Salt Lake City, USA; Division of Nephrology, Department of Internal Medicine, School of Medicine, University of Utah, Salt Lake City, USA; Medical Service, Veterans Affairs Salt Lake City Health Care System, Salt Lake City, USA; Department of Human Genetics, School of Medicine, University of Utah, Salt Lake City, USA; Department of Neurobiology, School of Medicine, University of Utah, Salt Lake City, USA

**Keywords:** *Drosophila*, ethanol, sensitivity, tolerance, genetics

## Abstract

Tolerance occurs when, following an initial experience with a substance, more of the substance is required subsequently to induce the same behavioral effects. Tolerance is historically not well-understood, and numerous researchers have turned to model organisms, particularly *Drosophila melanogaster*, to unravel its mechanisms. Flies have high translational relevance for human alcohol responses, and there is substantial overlap in disease-causing genes between flies and humans, including those associated with Alcohol Use Disorder. Numerous *Drosophila* tolerance mutants have been described; however, approaches used to identify and characterize these mutants have varied across time and between labs and have mostly disregarded any impact of initial resistance/sensitivity to ethanol on subsequent tolerance development. Here, we have analyzed a large amount of data – our own published and unpublished data and data published by other labs – to uncover an inverse correlation between initial ethanol resistance and tolerance phenotypes. This inverse correlation suggests that initial resistance phenotypes can explain many ‘perceived’ tolerance phenotypes. Additionally, we show that tolerance should be measured as a relative increase in time to sedation between an initial and second exposure rather than an absolute change in time to sedation. Finally, based on our analysis, we provide a method for using a linear regression equation to assess the residuals of potential tolerance mutants. We show that these residuals provide predictive insight into the likelihood of a mutant being a ‘true’ tolerance mutant, and we offer a framework for understanding the relationship between initial resistance and tolerance.

## 1. Introduction

Alcohol Use Disorder (AUD) is a major public health and societal problem. According to the 2021 National Survey on Drug Use and Health, 60 million people aged 12 years or older were binge drinkers in the past month, and 29.5 million people 12 years and older reported having a diagnosed AUD in the past year. Two main risk factors for AUD are resistance to the initial intoxicating effects of alcohol and elevated alcohol tolerance. Tolerance to ethanol occurs when, after an initial experience with the substance, a higher dose is required subsequently to induce the same behavioral effect. This phenomenon, also known as functional tolerance, is due to neuroadaptations,^1, 2^ but the specific neurobiological mechanisms underlying tolerance are still poorly understood. Indeed, tolerance is understudied, despite being one of the criteria for diagnosing AUD and critical to the persistence of alcohol abuse.^3^ A better understanding of the mechanisms of ethanol tolerance may lead to improved diagnosis and treatment of AUD.

*Drosophila melanogaster* is a proven useful model for studying the neurobiological and behavioral effects of alcohol. Indeed, flies display many of the behavioral responses observed in mammals, including humans, such as hyperactivity when exposed to low doses of alcohol and sedation with higher doses of alcohol. Flies, like humans, also show naïve avoidance of ethanol but learned preference upon repeated experiences, and they develop tolerance and experience withdrawal symptoms. Moreover, *Drosophila* share strong homology with human genes and conservation of signaling pathways and neurotransmitter systems, which allows genetic studies with high translational relevance for humans.^4–7^

Tolerance has been well-studied in *Drosophila*, and many tolerance mutants have been described (reviewed in ^5, 8^). However, methods used to determine tolerance mutants have varied. For example, tolerance was initially described using an inebriometer for ethanol exposure. This device measures the ethanol-induced loss of postural control where flies that become sedated elute out the bottom of the inebriometer, are counted, and cease to be exposed to ethanol. By quantifying the relative increase in the mean time to elution between the first and second exposure, tolerance is determined.^1^ In this approach, tolerance mutants are defined as flies with a significantly different percent increase in mean elution time from the genetic controls. It is worth noting that in the inebriometer, experimental and control flies may be exposed to a different dose of ethanol during the first, tolerance-inducing exposure. If experimental flies are resistant, for example, they may elute after 24 min of ethanol exposure, compared to 20 min for the control. Assuming similar ethanol absorption and metabolism (true for most published mutants), resistant flies will elute slower, be exposed to more ethanol, but be exposed to the same ‘effective dose’ i.e., the dose that causes complete loss of righting. However, other approaches to exposing flies to ethanol have been taken. Often, flies are exposed to ethanol vapor in closed containers which will not allow them to avoid exposure.^9^ In such setups, unlike in the inebriometer, flies continue to be exposed to alcohol even after becoming sedated. Thus, experimental and control flies can be exposed for the same duration (i.e., dose), allowing for a ‘fairer’ comparison of tolerance; after all, the development of tolerance is dose-dependent.^1, 10^

Previous research has sought to identify relationships between alcohol phenotypes. While ethanol tolerance and preference were found to be correlated, there was no relationship evident between initial resistance and tolerance.^11^ However, this was done in a fairly small sample. To determine whether the development of alcohol tolerance in any way depends on the initial resistance to the first sedation when the exposure dose is constant, we analyzed our own, and reanalyzed published tolerance data. We identified a highly significant inverse correlation between initial resistance (time to sedation) and tolerance (% increase in time to sedation). Our findings suggest that numerous genetic manipulations that, at face value, might be considered tolerance mutants may in fact be misclassified because they show initial sedation-sensitivity or sedation-resistance, which affects the development of tolerance secondarily. Knowing this inverse correlation also suggests that ‘true’ tolerance mutants are determined by correcting for this correlation, and we propose a predictive correlation function to aid in identifying *Drosophila* mutants that specifically affect the development of ethanol tolerance.

## 2. Materials and Methods

### 2.1 Fly stocks and genetics

Fly rearing and crosses were done on a 12:12-h light-dark cycle on regular cornmeal/yeast/molasses food at 25°C/65% humidity (unless otherwise specified). *w*Berlin* flies were used as +/+ controls. Some defined mutant alleles flies were outcrossed for 5 generations to the *w*Berlin* genetic background. Other experimental flies (e.g., *UAS-RNAi* lines for knock- down) were compared to siblings that lacked the *UAS-RNAi* transgenic insert or to control flies that served to inject DNA to generate the *UAS-RNAi* line. Many fly strains were obtained from the Bloomington or Vienna stock centers or were generous gifts from colleagues. All fly lines used in this study are listed in *Table 1*.

**Table 1:**
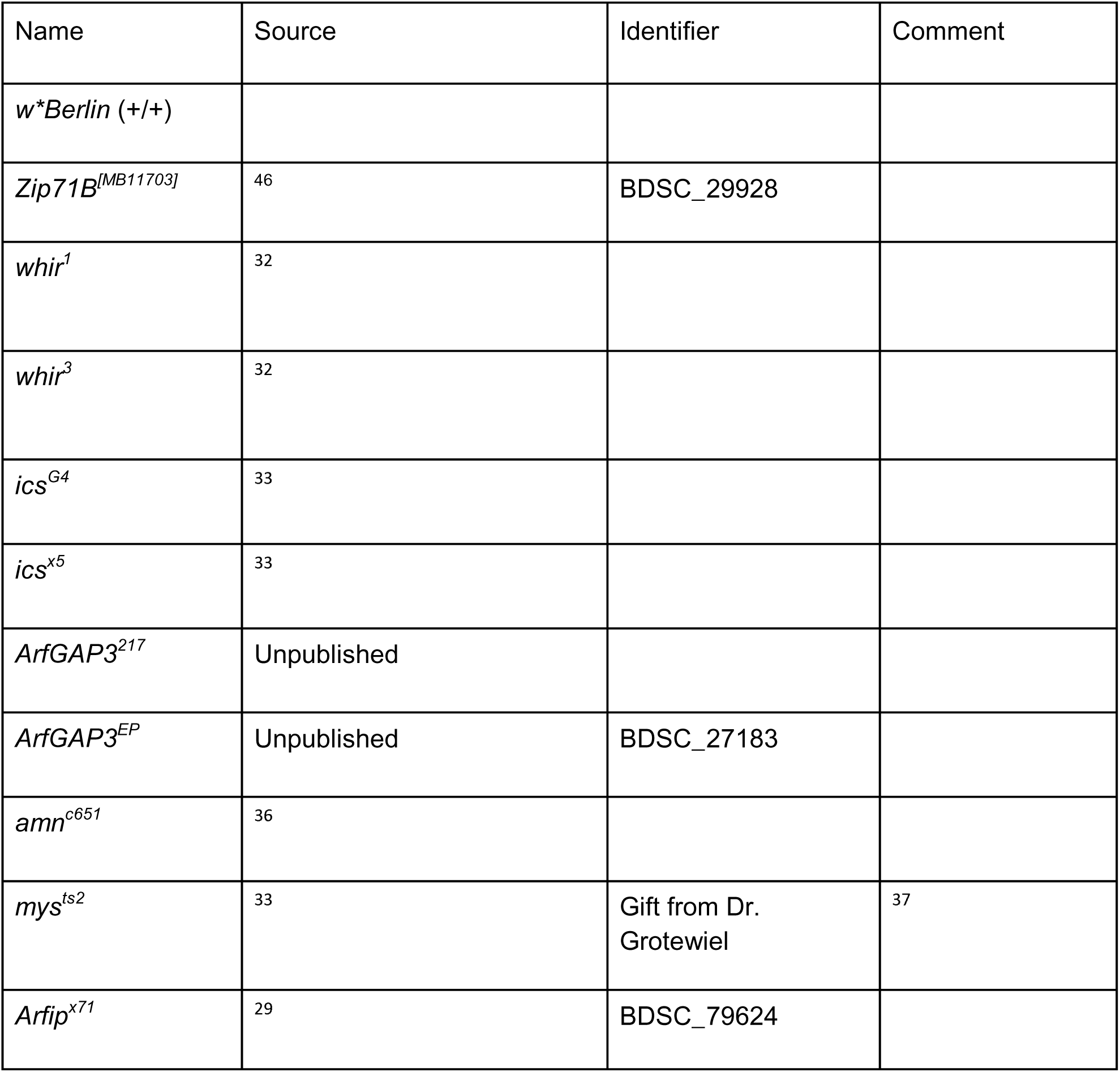
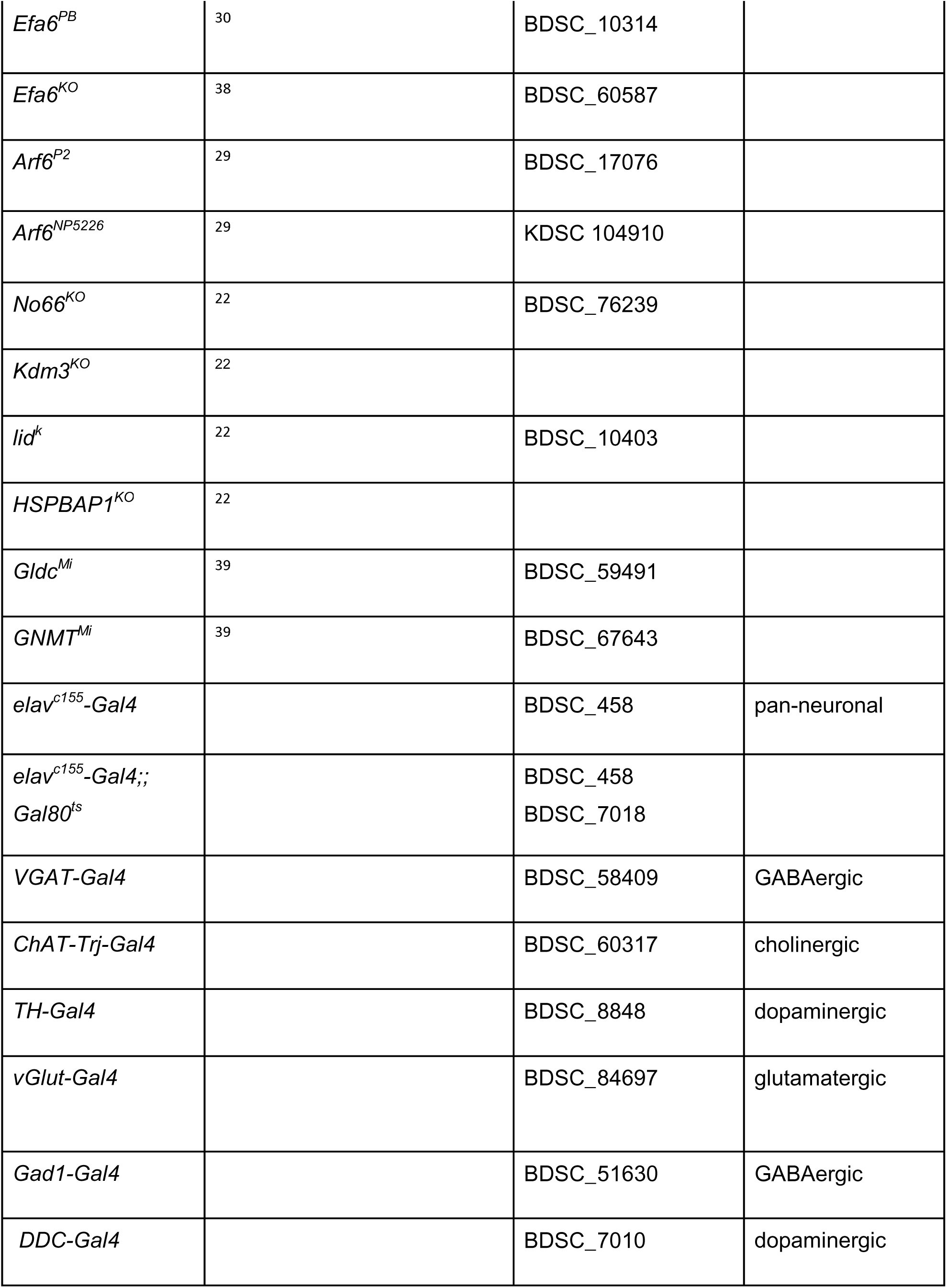
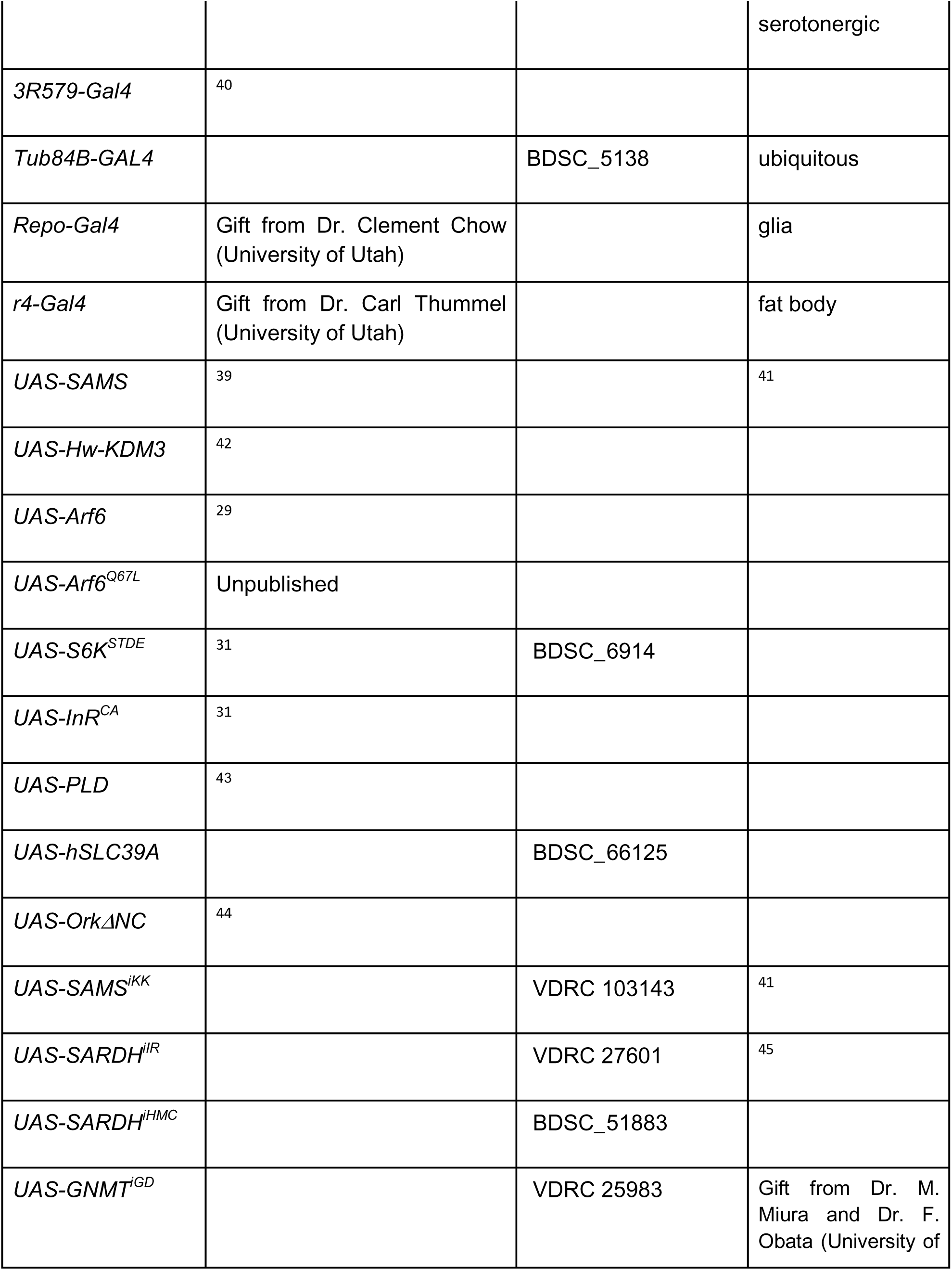
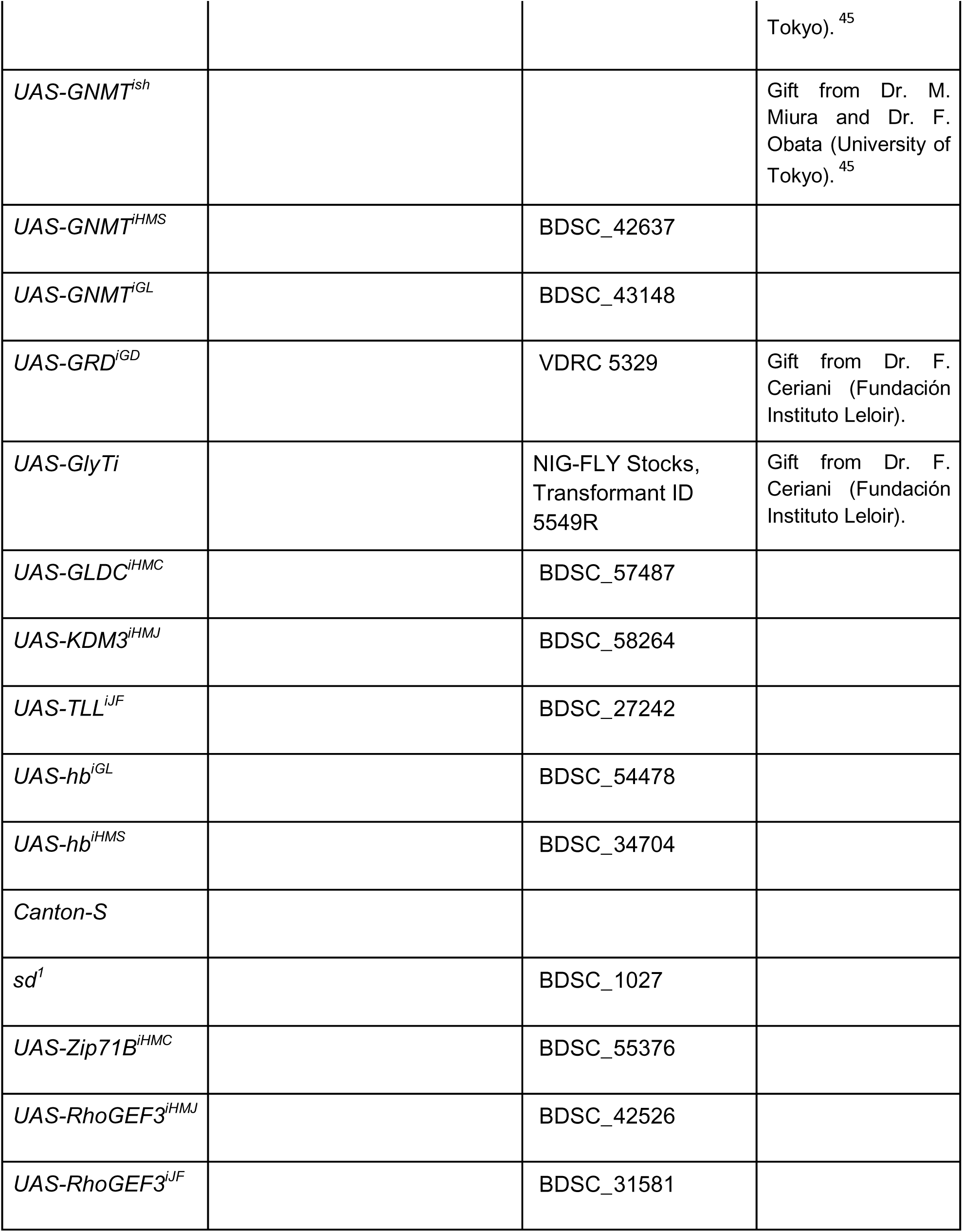
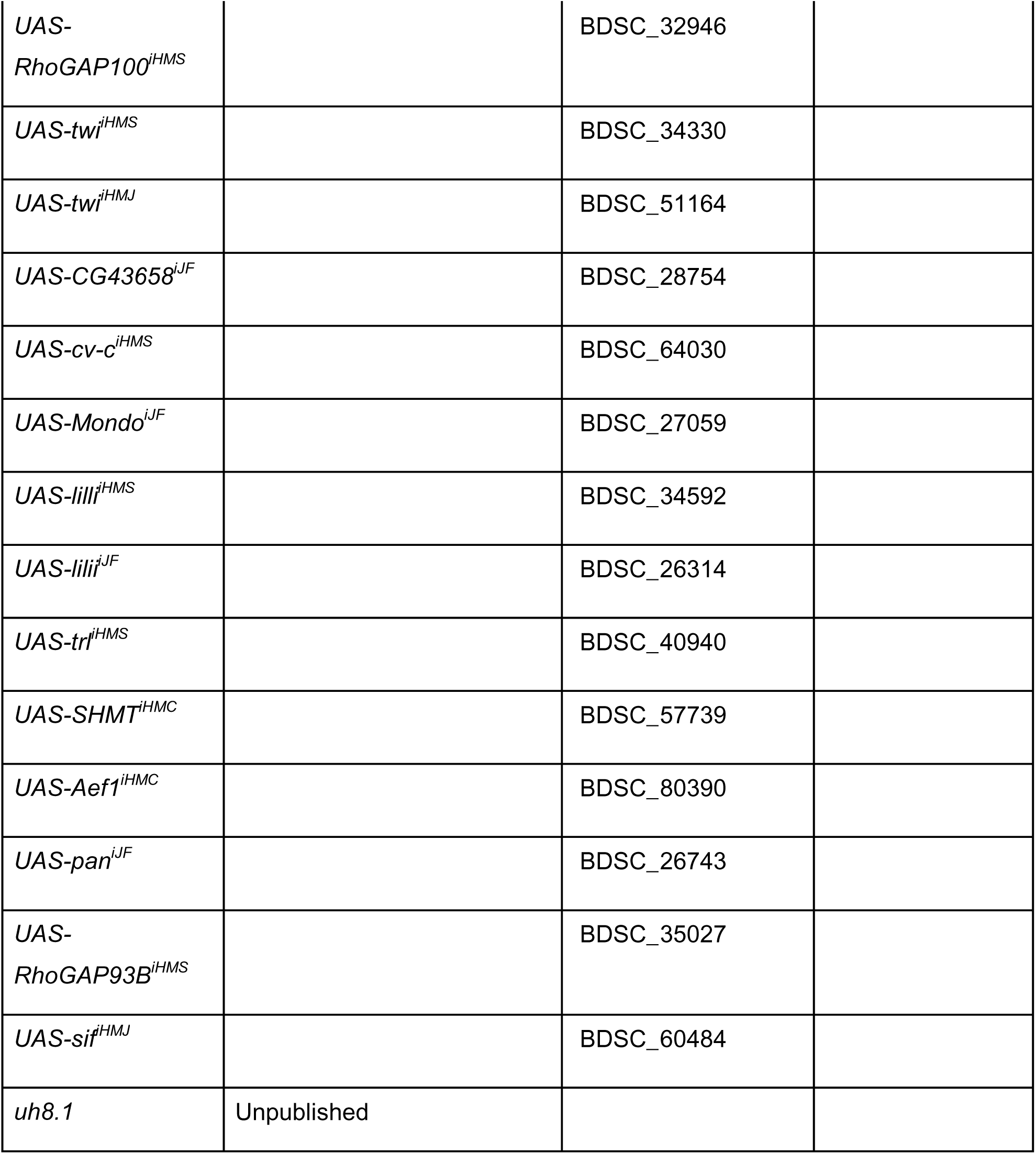
List of fly lines used in our laboratory for this study.

### 2.2 Behavioral experiments

3-7 day old adult males were collected after eclosion and used for experiments after at least 16 hours of recovery. Ethanol exposure and determination of ST-50 (time to half the flies being sedated) via measuring the flies’ loss-of-righting reflex (LORR) was performed as described previously.^12^ Briefly, flies were exposed to ethanol vapor and visually inspected every 2 minutes for LORR after light tapping, and the ST-50 for 8-10 flies was determined and counted for an n of 1. For tolerance, flies were exposed to ethanol for 22 min during first exposure and re-exposed 4 hours later. Each set of experimental and control flies was assayed in parallel on the same day and repeated at least 2 times on different days. For Figure 4 and Figure 6, flies were exposed to 1.5 times their ST-50, which was individually determined for each n of 1 (8-10 flies). Tolerance was calculated as the % increase in ST-50 from exposure 1 to exposure 2.

### 2.3 Ethanol absorption

Ethanol concentration in flies was measured using the method established by Ishmayana et al.^13^ Control and experimental flies (a total of n=10 per condition were tested where n=1 consisted of 10 flies) were exposed to ethanol vapor for 1.5 times the ST-50. At the end of the exposure, flies were frozen and homogenized. 2µl of homogenates were used to measure ethanol concentration at 340nm on a Nanodrop. Reagents used for this assay were Alcohol Dehydrogenase (ThermoFisher, Cat No J65869.9+, lot S301038) and beta nicotinamide adenine nucleotide (ThermoFisher, Cat No J62337.03+, lot M251003).

### 2.4 Experimental Design and Statistical Analyses

Data are represented by mean +/- SEM for ethanol sedation and tolerance data. Data were analyzed using GraphPad Prism Software version 9, and statistical analysis was done with unpaired Student’s t-test. We then calculated the effect sizes with Hedge’s *g* for ST-50 and %Tolerance (± 95% confidence intervals for some panels). We plotted the effect size for ST-50 on the x-axis and the tolerance effect size on the y-axis and performed a simple linear regression. We also determined ST-50s and tolerance from previous published studies.^14–21^ Our inclusion criterion was that the control and experimental flies had to be subjected to the same dose of ethanol for the same period of time. For previously published data, we included 2 experimental manipulations per gene, generally using a classical mutant and a knockdown in specific regions when available.

## 3. Results

### 3.1 Inverse correlation of initial resistance and tolerance phenotypes in our mutants

To test for a relationship between the resistance to naïve alcohol-induced sedation (henceforth referred to as initial resistance) and alcohol-induced tolerance, we have previously examined a family of jumonji-domain histone demethylase (*HDM*) mutants and found no correlation.^22^ We re-examined these data and calculated the effect sizes compared to wild-type for ethanol resistance, as measured by the ST-50, the time to sedation for half of the flies. An effect size > 0 signifies increased initial resistance (Figure 1A). We also determined effect sizes for tolerance, as measured by the percentage increase in ST-50 from the first to the second exposure. As before, an effect size > 0 signifies the development of more tolerance (Figure 1A). In these experiments, experimental genotypes and control flies were exposed to alcohol for the same duration of time. There was no correlation between effects on initial resistance and tolerance for the whole cohort of 13 loss-of-function (LOF) *HDM* mutants (Figure 1B). However, when we focused on the 6 *HDM* mutants that showed a significant change from the wild-type control in initial resistance and/or tolerance, we noticed that 5 of the 6 mutants lay in effect size quadrants where increased initial resistance correlates with decreased tolerance and vice versa – the exception being *Kdm3^KO^* mutants, which show less initial resistance and develop less tolerance (Figure 1B). For 4 of these 6 mutants, including Kdm3, we also determined the responses to different alcohol exposure doses (ranging from high to low EtOH concentrations), and the 3 non-*Kdm3* mutants again showed an inverse correlation between initial resistance and tolerance (*R*^2^=0.63, F(1,12)=20, *p*=0.0008; Figure 1C). Even including Kdm3, the correlation was significant (*R*^2^=0.25, F(1,17)=5.6, *p*=0.031).

**Figure 1.**
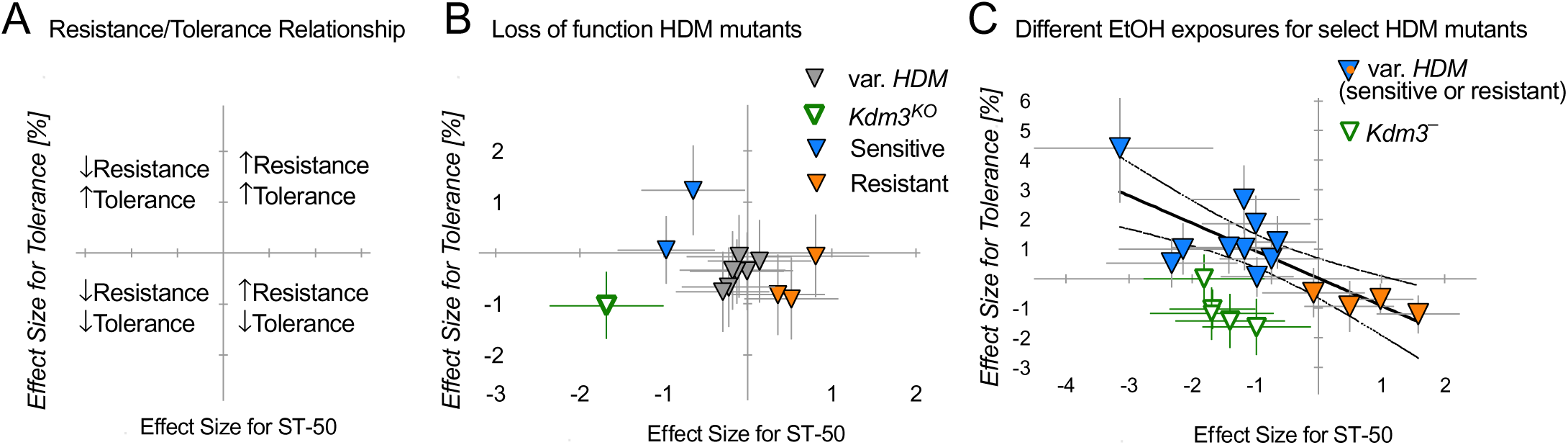
**A** - Plot of effect sizes for experimental vs. control phenotypes of sedation (ST-50) on x-axis and effect sizes for tolerance on y-axis. Data points in the upper left quadrant show decreased initial resistance during the first exposure with increased tolerance after the second exposure, decreased resistance and decreased tolerance for lower left quadrant, increased resistance and decreased tolerance for the lower right quadrant and increased resistance with increased tolerance for the upper right quadrant. **B** - Effect sizes for tolerance and for sedation for loss-of-function histone demethylase (HDM) mutants were plotted. Gray symbols depict HDM mutants with no resistance or tolerance phenotype. Blue symbols depict initially sensitive mutants, orange symbols resistant mutants, and green is a *Kdm3^KO^* mutant, a possible ‘outlier’.^22^ Here, and in subsequent panels, effect size error bars are 95% confidence intervals. **C** - Effect sizes for additional tolerance and sedation assays for loss-of-function HDM mutant that showed a phenotype previously (in B). These mutants were exposed to different alcohol concentrations.^22^ There is a significant inverse correlation between initial resistance and tolerance with these mutants when *Kdm3* is excluded. The linear regression plotted was calculated excluding *Kdm3* data points; however, even with *Kdm3* data included, there is a significant correlation (see text for details). Mutants include knock outs and neuron-specific knock-down by RNAi.^22^

To determine whether there was indeed an inverse correlation between initial resistance and tolerance phenotypes, we examined additional genetic manipulations that caused significant initial resistance and/or tolerance phenotypes when experimental and control flies were exposed to the same dose of alcohol. First, we analyzed loss-of-function (LOF) mutants in 12 non-*HDM* genes we have studied over the years (most published, but some unpublished), and 10 showed an inverse correlation between resistance and tolerance effect sizes (*R^2^*=0.80, F(1,13)=52, *p*<0.0001; Figure 2A). The two exceptions were mutants in *Arf6* and its activator *Efa6,* which both showed reduced initial resistance and reduced tolerance (Figure 2A), similar to *Kdm3* (Figure 1B). Although, the correlation was significant even including Arf6 and Efa6 (*R^2^*=0.32, F(1,17)=8.1,*p*=0.011). Including the above *HDM* mutants, of the 16 genes examined, 13 LOF mutants showed inversely correlated effects on initial resistance and tolerance.

**Figure 2.**
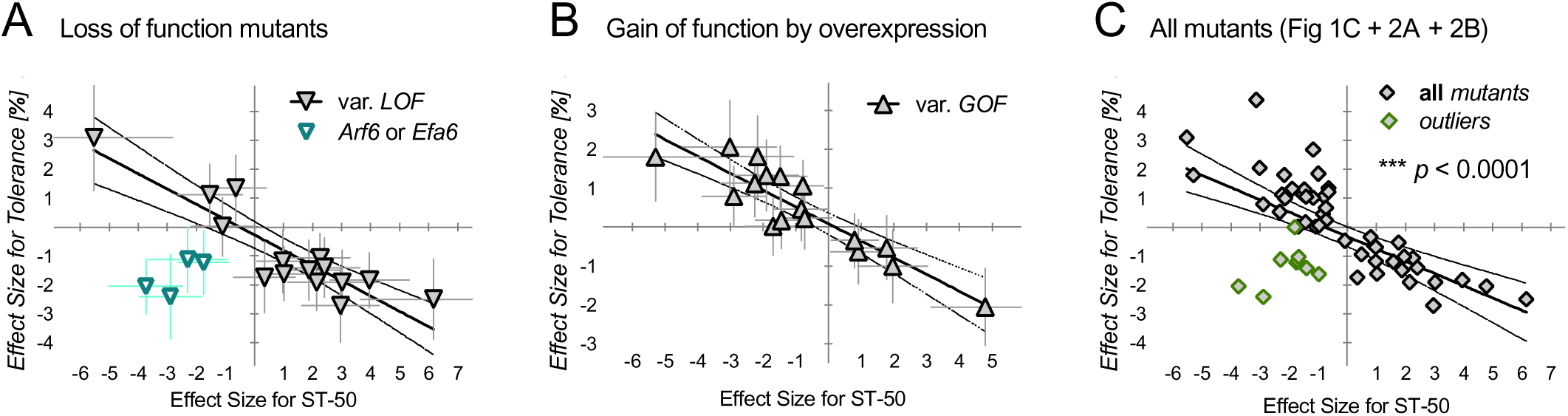
**A** - Effect sizes for tolerance and for sedation for various of loss-of-functions (LOF) mutants from our lab, including *Arf6* and *Efa6* mutants, were plotted (^29, 33, 39, 46,unpublished^ ^data^). Teal symbols depict *Arf6* and *Efa6* mutants, who are putative outliers. The linear regression was calculated excluding Arf6 and Efa6 data points, but inclusion of these mutants still resulted in a significant correlation (see text). **B** - Effect sizes for tolerance and for sedation for various gain-of-function (GOF) neuronal manipulations were plotted. The linear regression was calculated including all data points. **C** - Effect sizes for tolerance and for sedation for all of our mutants (data from Figure 1C, 2A and 2B) were plotted. The linear regression was calculated including all data points. Putative outliers (*Kdm3*, *Arf6*, *Efa6*) are depicted by a green symbol, but are included in the correlation calculation.

We also analyzed several gain-of-function (GOF) mutants, using the *Gal4/UAS* system to overexpress wild-type or constitutive-active versions of various genes in a neuron-specific manner (i.e., in all neurons or in subsets of neurons such as GABAergic or cholinergic neurons). We analyzed 7 genes in 17 experiments, using 9 different *Gal4* drivers, which also showed a significant inverse correlation between their GOF initial resistance and tolerance phenotypes (*R^2^*=0.81, F(1,15)=65, *p*<0.0001; Figure 2B). Taken together, the combined 55 experiments shown in Figures 1 and 2, manipulating 21 different genes, showed an inverse correlation with an *R^2^* of 0.4 and *p* < 0.0001 (F(1,53)=35; data from Figures 1C, 2A, and 2B, no ‘outliers’ removed and presented together in Figure 2C). When we removed the putative outliers, *Arf6*, *Efa6*, and *Kdm3* (9 experiments), the remaining 46 experiments showed an inverse correlation with *R^2^*= 0.74 (F(1,44); *p* < 0.0001; Figure 2C). Thus, when experimental and control flies are exposed to the same dose of alcohol in our hands, effects on initial resistance and tolerance are inversely correlated.

### 3.2 Inverse correlation of initial resistance and tolerance phenotypes in other published mutants

Since all data in Figures 1 and 2 were generated in our own lab, we wondered whether our observed inverse correlation would hold true for published phenotypes from other labs. We combed the literature and analyzed publications from the last 15 years that determined initial resistance and tolerance in ways similar to ours, i.e., parallel exposure of experimental and control flies to the same dose of ethanol. This left us with 8 papers from 6 labs describing 28 manipulations in 16 genes and their resultant ethanol-related phenotypes. We analyzed published mutants with significant initial resistance and/or tolerance phenotypes, again revealing a significant inverse correlation between initial resistance and tolerance (Figure 3A; *R^2^* = 0.57, F(1,26)=35, *p* < 0.0001). Together, these analyses illustrate that in our hands and other labs, there is a significant inverse correlation between initial resistance and relative tolerance when experimental and control flies are exposed to the same dose of alcohol.

**Figure 3.**
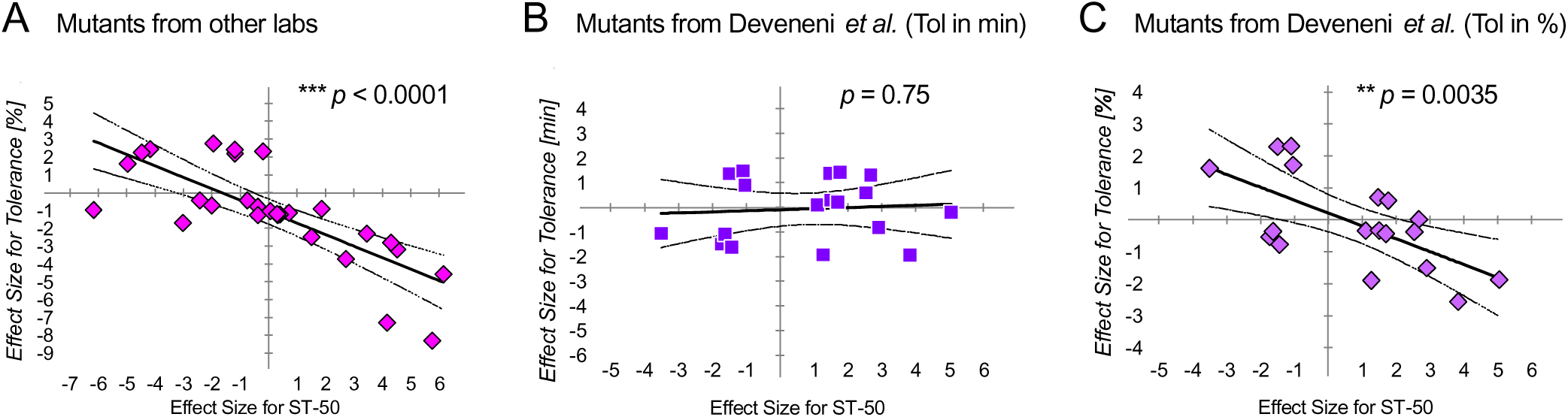
**A** - Effect sizes for tolerance and for sedation for various mutants published by other labs.^14–16, 18–21, 26^ The linear regression was calculated including all data points. **B** - Effect sizes for tolerance and for sedation for mutants published in Devineni et al.,^11^ where tolerance is calculated in absolute terms, i.e., difference in minutes. The linear regression was calculated including all data points. **C** - Effect sizes for tolerance and for sedation for mutants shown in B, where tolerance is re-calculated in relative terms, i.e., % increase in ST-50. The linear regression was calculated including all data points.

Devineni and colleagues ^11^ had also previously asked whether initial resistance and tolerance phenotypes correlate, analyzing 18 mutants in various genes from a large forward genetic screen for alcohol-response phenotypes. Unlike in our analysis, where tolerance was calculated as relative (percent increase from exposure 1 to exposure 2), they measured tolerance as an absolute increase in minutes from exposure 1 to exposure 2. In doing this, they found no correlation between initial resistance and tolerance. This data, replotted for effect sizes, is shown in Figure 3B (*R^2^* = 0.006, F(1,16)=0.1; *p* = 0.75; not significant, as published by ^11^). Using these data, we re-calculated tolerance as a percent increase relative to the initial resistance measured by the initial ST-50 (as in Figures 1 and 2) and plotted the resulting effect sizes. When measured this way, there was again a significant inverse correlation between initial resistance and tolerance (Figure 3C; *R^2^* = 0.42, F(1,16)=12, *p* = 0.0035).

### 3.3 Tolerance should be expressed in relative, not absolute, terms

As outlined above, we determined a significant inverse correlation between initial resistance and tolerance phenotypes when tolerance is expressed as a fractional change relative to the first ST-50. While some other labs also calculate tolerance this way, other publications, including the one described in Figure 3B-C,^11^ express tolerance as an absolute difference in minutes, subtracting the initial ST-50 from the second ST-50.^23^ Since the inverse correlation that we found holds true for tolerance analysis in relative terms (see Figure 3B vs. 3C), we wanted to determine which of the two measures is more accurate for analyzing tolerance.

Increasing the dose (i.e., duration) of the first ethanol exposure increases tolerance.^1^ This increase in tolerance is reflected in both relative and absolute terms and is therefore not helpful in determining the best way to calculate tolerance. However, we reasoned that when flies are exposed to the same initial dose of ethanol, tolerance should be the same. We therefore exposed flies of the same genotype (*w*Berlin*) to two different concentrations of ethanol (95% vs. 80%), where flies receiving the higher concentration were exposed for a shorter time than flies receiving the lower concentration (Figure 4A). In both cases, flies were exposed to 1.5 times the ST-50, which we independently determined for each vial of flies tested, resulting in complete loss of the righting reflex. After both the first and second exposure, the ST-50 was significantly shorter in the 95% EtOH-exposed flies (8.9 minutes on the first exposure and 11.5 minutes on the second exposure) than in the 80% EtOH-exposed flies (12.4 minutes on the first exposure and 16.4 minutes on the second exposure; Figure 4B). Importantly, these two exposure paradigms resulted in indistinguishable internal ethanol concentrations after the first exposure (t(18)=0.6, *p*=0.55; Figure 4C), suggesting that these flies indeed received the same exposure doses. We then quantified the tolerance these two exposures induced in both relative and absolute terms. The amount of tolerance was significantly different when calculated in absolute terms as minutes (*p*=0.040, t(22)=2.2; Figure 4D), while it was not significantly different when expressed in relative terms, as percent increase (*p*=0.40, t(22)=0.4; Figure 4E). These data show that our two paradigms of exposing flies to ethanol resulted in the same initial exposure dose (as measured by internal ethanol concentration after exposure 1). Behaviorally, they gained the same amount of tolerance in percent, while tolerance expressed in absolute minutes was significantly different. Given that the same genotype of flies received the same dose of ethanol in both exposures, our data indicate that ethanol’s effect on tolerance should be expressed fractionally relative to the initial resistance. These data also support our finding of an inverse correlation between initial resistance and tolerance phenotypes.

### 3.4 Additional evidence to support an inverse correlation

Given our determination that expressing tolerance relative to initial resistance is an accurate measure and that there is a significant inverse correlation between these two, we were interested in expanding our analysis. In addition to examining various Mendelian mutants (Figures 1-3), we have also used RNA-interference (RNAi) to knock down the expression of many genes. We collated all the data for these experiments, again analyzing experiments run in parallel where experimental flies showed a significant difference in initial resistance and/or tolerance from the controls. Here, we note that we have not verified the knock-down efficiency of the RNAi transgene for most of these experiments, nor have we ruled out any possible off-target effects. We, therefore, do not claim that the presumed target genes are responsible for the observed phenotypes. However, we stress that experimental and control flies were exposed simultaneously, in parallel, to identical ethanol concentrations and durations and that they are genetically distinct from each other and showed significant resistance and/or tolerance phenotypes. Therefore, these experiments are well-suited to answer whether genetic differences cause correlated ethanol response phenotypes in initial resistance and tolerance. We found that in 52 additional experiments, with 32 distinct RNAi-transgenes (putatively targeting 23 genes), using 10 different *Gal4* drivers, there was also a significant inverse correlation between initial resistance and tolerance (*R^2^* = 0.78, F(1,51)=180, *p* < 0.0001; Figure 5A). We combined the analyses from Figures 2C, 3A, 3C, and 5A into a single plot (Figure 5B) which shows a significant inverse correlation across all experiments and manipulations (*R^2^* = 0.51, F(1,157)=162, *p* < 0.0001; no ‘outliers’ excluded).

**Figure 4.**
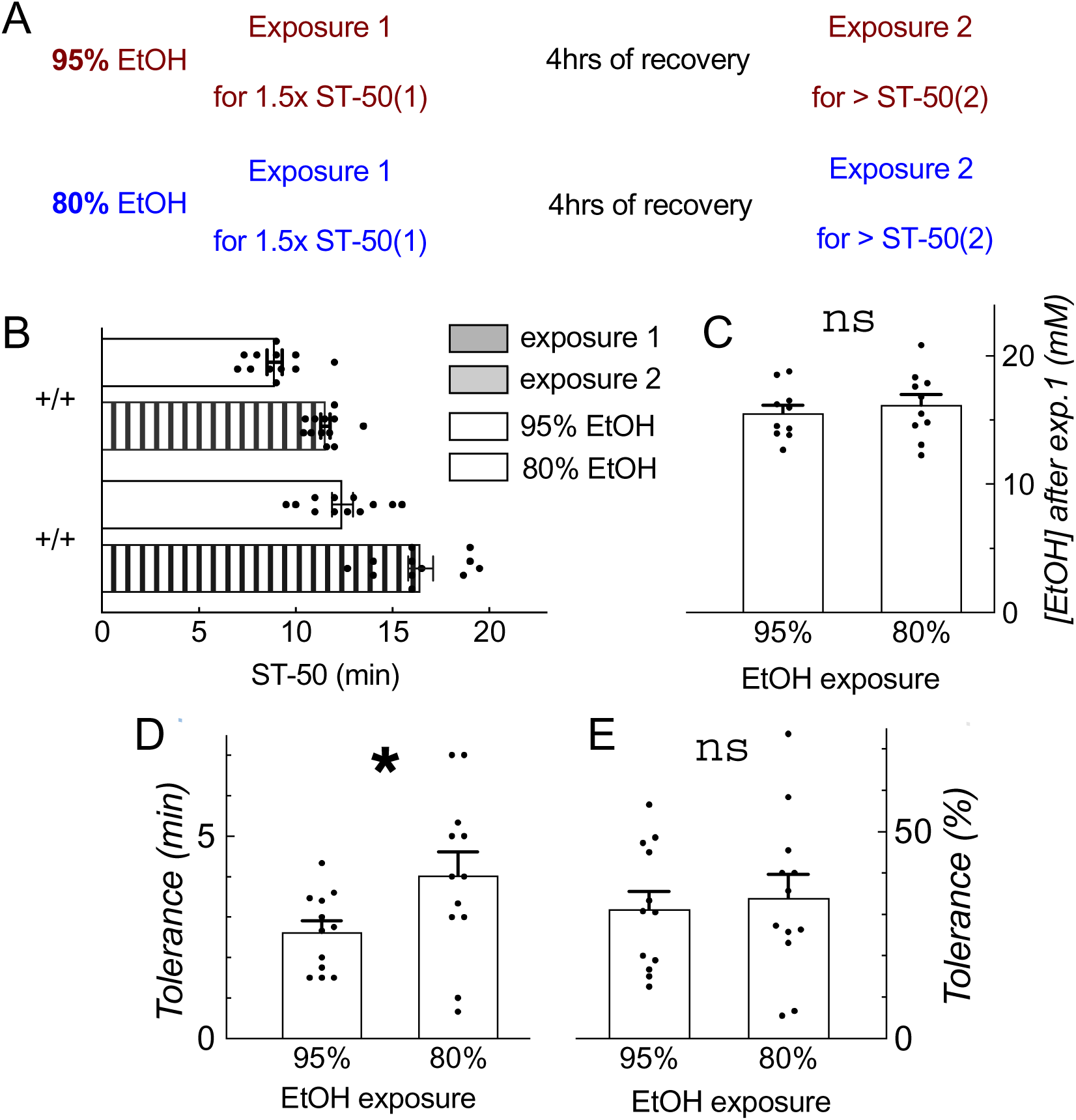
**A** - Schematic of EtOH exposure paradigms to determine if tolerance should be calculated in minutes or % increase. The same genotype, *w*Berlin*, was exposed in the two different ways shown. **B** - ST-50s of first and second exposures for flies exposed to 95% EtOH (brown) and 80% EtOH (blue), *n*=12 each. Data in this figure are shown as mean ±SEM. **C** - Internal ethanol concentrations (mM) in flies collected after the first exposure to either 95% or 80% EtOH. **D, E** - Tolerance developed in flies exposed to either 95% or 80% EtOH expressed in absolute terms (D, difference in minutes) and in relative terms (E, % increase).

**Figure 5.**
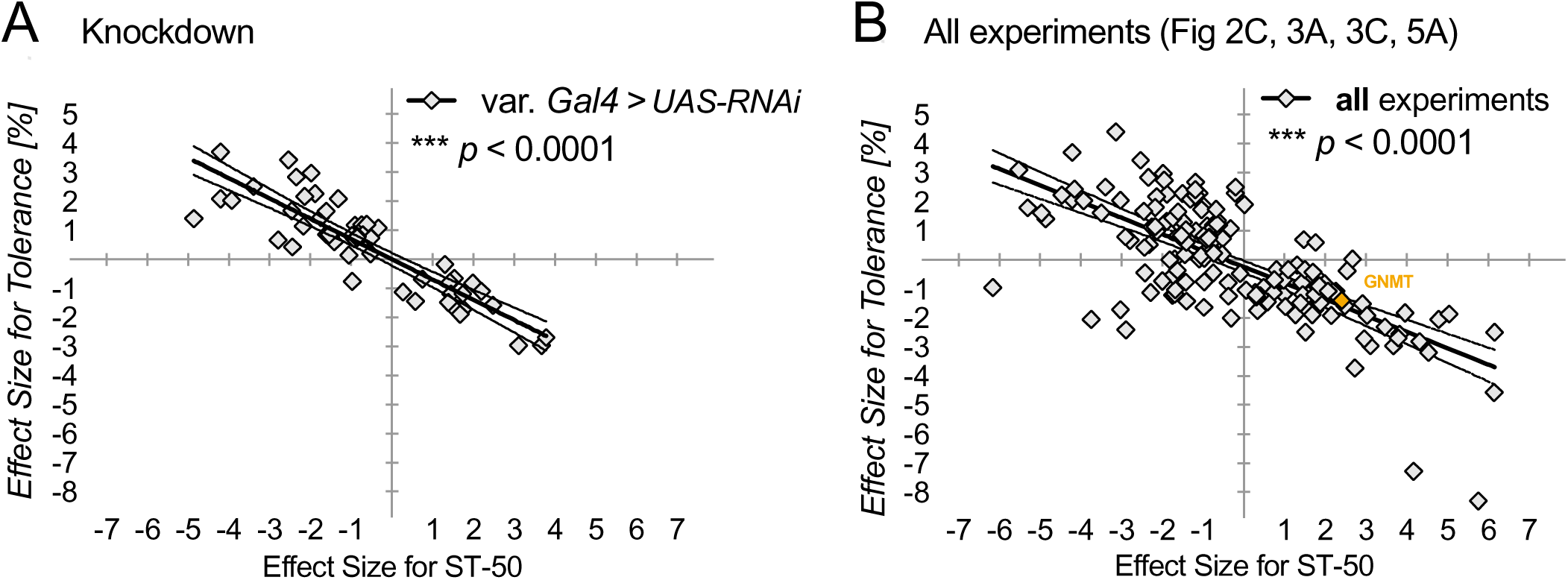
**A** - Effect sizes for tolerance and for sedation for various Gal4-mediated overexpression and knockdown experiments performed in our lab. The linear regression was calculated including all data points. **B** - Effect sizes for tolerance and for sedation for all mutants analyzed (data from Figure 2C, 3A, 3C and 5A) The linear regression was calculated including all data points with. *GNMT* mutant is highlighted in yellow.

### 3.5 Tolerance phenotypes can be a consequence of initial resistance phenotypes

Our inverse correlation shows that flies that are more sensitive to ethanol (i.e., become sedated quickly) generally develop more tolerance, and resistant flies less. One explanation for this is that highly sensitive flies that are exposed for the same duration as less-sensitive controls achieve a greater depth of sedation which facilitates the development of more tolerance. Therefore, it may be the case that the absolute dose in mM ethanol does not matter as much for the homeostatic process of tolerance but that it is the depth of sedation and, with it, the effectiveness of depression of CNS activity, that determines tolerance. Indeed, longer first ethanol exposures are known to induce more tolerance,^1, 10^ presumably because they depress CNS activity more strongly and for a longer duration.

What implications does this have when determining tolerance for a mutant that affects initial resistance? Since exposing a given mutant and its control to the same dose may result in a distinct depth of sedation/effectiveness of CNS depression, we reasoned that they should be exposed to the same ‘effectiveness of intoxication’, most easily measured by the behavioral effect on sedation. For example, in the inebriometer, flies are exposed to complete loss-of-righting, fall through the exposure column, and are no longer exposed. Thus, experimental and control flies are exposed to the same behavioral effect, or effectiveness of intoxication. If this complete loss-of-righting induced distinct amounts of tolerance, then such flies truly are tolerance mutants.

Conversely, mutants whose sedation and tolerance effects fall onto the inverse correlation curve only appear mutant due to distinct levels of effectiveness of intoxication during the first exposure. GNMT is an enzyme involved in the 1-Carbon cycle, and when exposed for the same duration as controls (Figure 6A), *GNMT* LOF causes increased initial resistance (t(21)=4.0, *p*=0.0007; Figure 6B) and reduced tolerance (t(20)=3.8, *p*=0.0012; Figure 6C). At face value, this data suggests that *GNMT* is a tolerance mutant. However, the ‘tolerance phenotype’ was exactly as predicted by the inverse resistance/tolerance correlation (see Figure 5B). We therefore predicted that if we exposed wild-type and *GNMT* mutants to the same depth of sedation – 1.5 times their respective initial ST-50 – they would both develop the same amount of tolerance. Indeed, exposing *GNMT* mutants and wild-type to ethanol for 1.5*ST-50 necessitated longer initial exposures in *GNMT* mutants (15.5 minutes vs. 12 minutes for wild-type; Figure 6D). Although there was a significant difference in initial resistance (t(21)=4.2, *p*=0.0004 Figure 6E), there was no difference in the subsequent development of tolerance (t(21)=1.3, *p*=0.20; Figure 6F) using this exposure paradigm. Therefore, *GNMT* may at first appear to be a tolerance mutant due to its significantly lower development of tolerance when exposed to the same dose as wild type, but we suggest that in that exposure paradigm, *GNMT’s* tolerance effects are simply a consequence of the initial resistance. Indeed, when *GNMT* mutants and wild-type flies receive an amount of alcohol that induces equivalent behavioral effects, the tolerance phenotype is lost. Knowing the inverse correlation of alcohol resistance and tolerance, therefore alters the interpretation of the initial *GNMT* tolerance phenotype.

## 4. Discussion

Here, we analyzed a large collection of genetic manipulations that result in ethanol resistance and/or tolerance phenotypes and detected a significant inverse correlation of the two measures when both experimental and control flies were exposed to the same initial, tolerance-inducing ethanol dose. Mutants with increased initial resistance developed less tolerance, and ones with decreased initial resistance developed more tolerance. One possible explanation of this inverse correlation is that apparent tolerance phenotypes arise in more sensitive mutants because they become more deeply sedated (and for a longer duration) after the first exposure, and they therefore developing more tolerance. This interpretation is consistent with published findings showing that larger exposure doses induce more tolerance.^1, 10^ Conversely, mutants that are resistant to ethanol are only lightly sedated at the end of the exposure compared to controls and therefore develop less tolerance. The inverse correlation may reflect the homeostatic control of CNS activity. If tolerance is caused by upregulation of neuronal activity triggered by prior ethanol-induced depression of neuronal activity,^8^ then the extent of tolerance developed may reflect how effective/deep the initial ethanol-induced neuronal depression was for a given first ethanol dose. This would support the idea that the functional effect of ethanol on the neurons matters more than the actual dose of ethanol. Thus, when the tolerance-inducing mechanism is intact, slight neuronal depression in resistant mutants and strong neuronal depression in sensitive mutants will induce slight or strong tolerance, respectively. This relationship would result in our observed inverse correlation and suggests that mutants develop tolerance as a function of their initial resistance. Therefore, ‘true’ tolerance mutants will show resistance and tolerance effects that do not lie on the inverse correlation curve but will instead be far away from that curve.

The distance from the inverse correlation curve is expressed by the residuals (Figure 7A). Figure 7B shows the distribution of residuals from the correlation, which could be used as a guide to determine which mutants are indeed tolerance mutants and which ones to follow up on. These residuals can be considered ‘corrected’ effect sizes for the tolerance phenotypes. Classically, effect sizes of ±0.8 or higher/lower are considered strong, which would apply to 71 out of 159 tolerance residuals. For the initial resistance phenotypes, 127 out of 159 data points lie beyond ±0.8. This suggests that many more of our analyzed genotypes are strong initial resistance mutants than are strong tolerance mutants, consistent with our finding that tolerance is affected by initial resistance. This also brings up the question: where should the tolerance residual cutoff be to consider a mutant a ‘true’ tolerance mutant? This cutoff is in the eye of the beholder, and from our experience, we prefer to work with strong mutants that have effect sizes beyond ±1.2. This would hold true for 45 tolerance residuals. However, because tolerance data are more variable than initial resistance data, we prefer an even larger effect size of ±1.5, which is the case for 36 tolerance residual data points. Regardless of the exact threshold chosen, in Figure 6, we show that *GNMT* is not a ‘true’ tolerance mutant since its tolerance phenotype disappears when mutants are exposed to the same depth of sedation as controls. Consistent with this, the calculated residual for *GNMT* in the initial experiment is 0.18 (Figure 7A, C-D), indicating that this mutant develops tolerance as expected from the inverse correlation and is therefore not a tolerance mutant.

**Figure 6.**
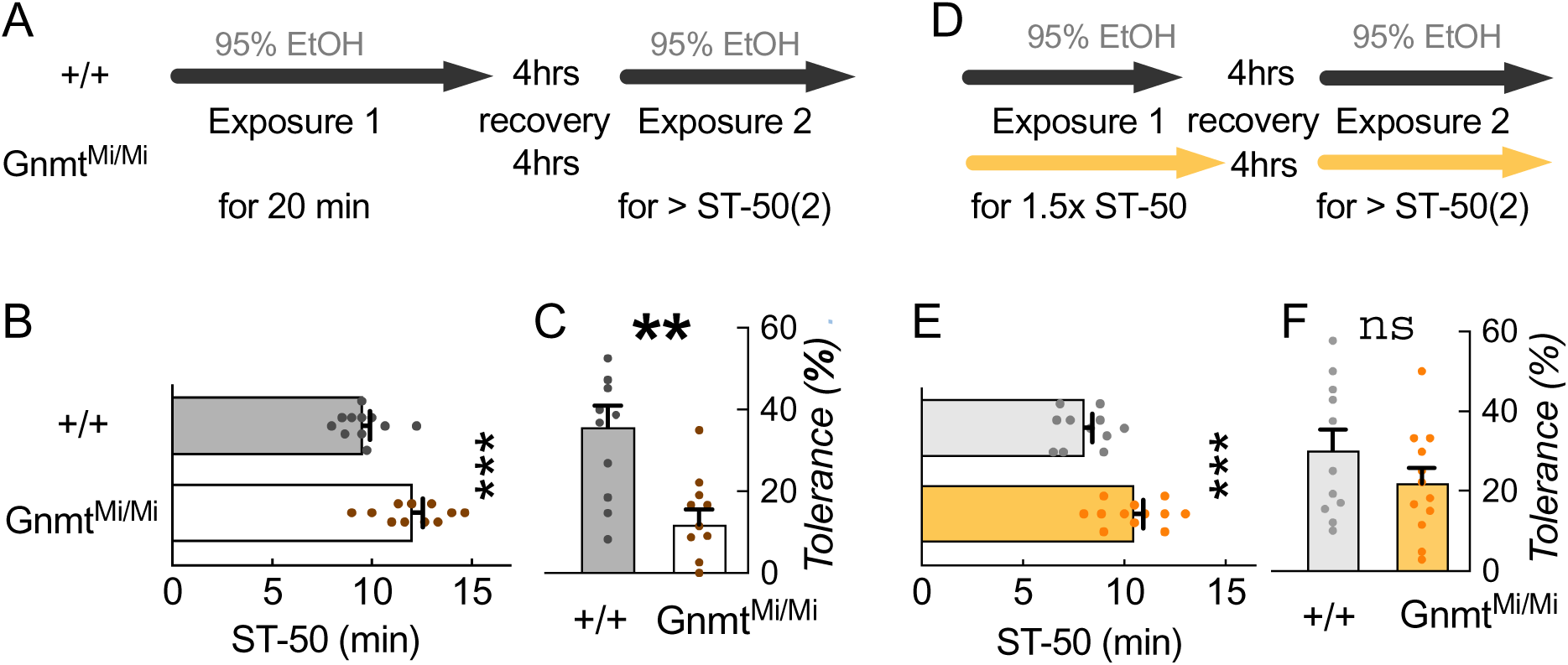
**A, D** - Schematic of EtOH exposure paradigm of *GNMT* mutant and control flies. **B, E** - Ethanol sedation phenotype for controls vs. *GNMT* mutants during regular exposure paradigm (A,B) or when exposed to 1.5*ST-50 (D,E). In both cases *GNMT* mutants are resistant to the initial ethanol-induced sedation. Data shown as mean ±SEM. **C, F** - Tolerance phenotype (% increase) for controls vs. *GNMT* mutants during regular exposure paradigm (C) or when exposed to 1.5*ST-50 (F).

**Figure 7.**
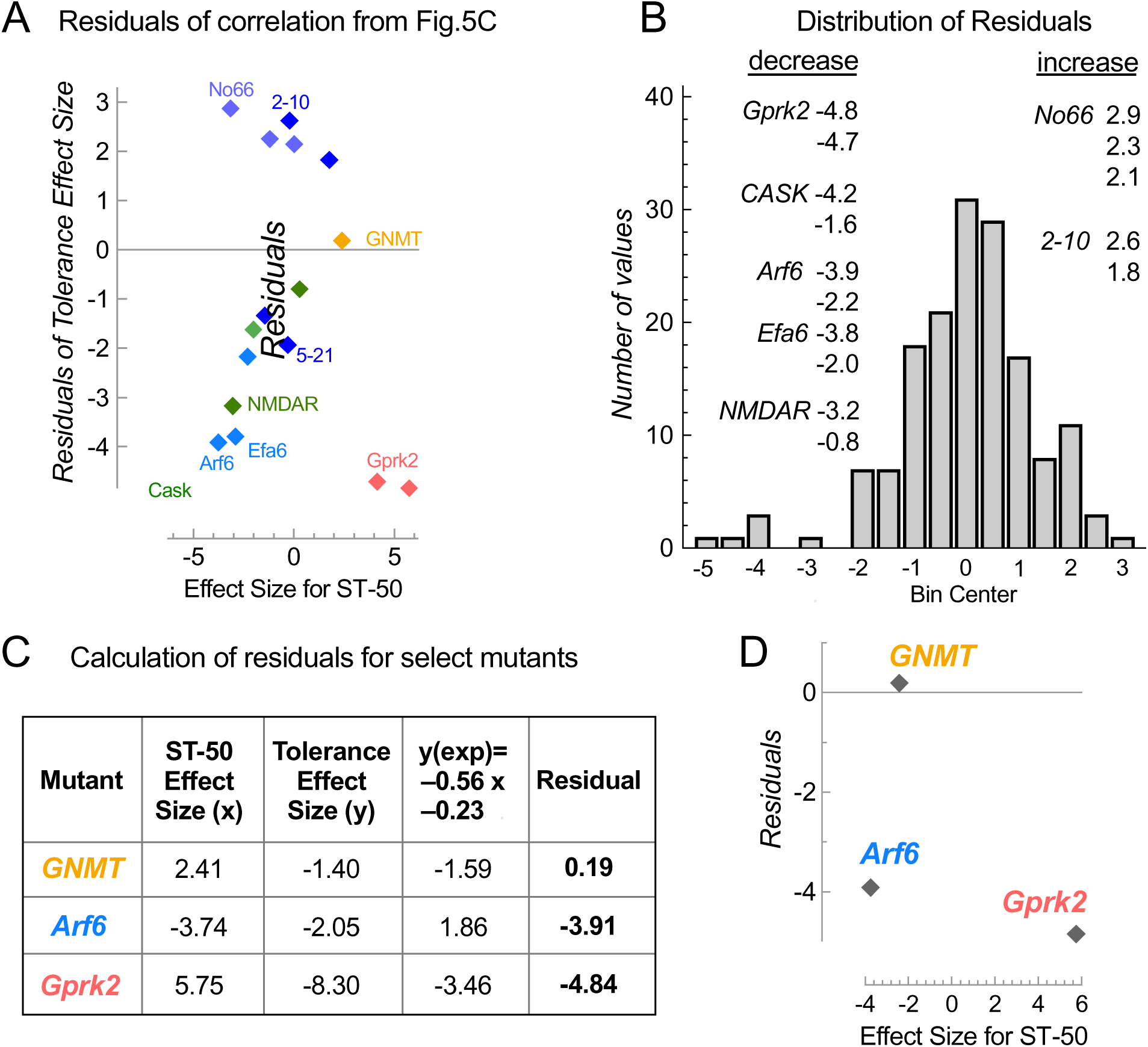
**A, B** – Residual plots for all the data analyzed (*n*=159). Residuals (y-axis) in the XY plot (A) indicate how far from the inverse correlation curve the data points lie, while the frequency plot. (B) shows a normal distribution of the residuals centered around 0 (mean=0.004). Some potential strong, true tolerance mutants are highlighted (*Arf6*, *Efa6*, *CASK*, *NMDAR*, *Gprk2*, *No66*, *2-10* and *5-21*) as well as *GNMT* (yellow), which is not a tolerance mutant. **C** - Example of how to calculate residuals using the equation y(exp) = –0.56 x –0.23 with *GNMT*, *Arf6* and *Gprk2* data, where y(exp) stands for the tolerance effect size (y) expected. **D** - Plot of residuals calculated in C versus effect size for ST-50.

The residual plots (Figure 7A, B) highlight a few strong ‘true’ tolerance mutants. Among the strong tolerance mutants are mutants *2-10* and *5-21* from Devineni *et al.*^11^ In 1 of 2 experiments, these showed the same initial ST-50 as the control and then developed more or less tolerance, respectively. By definition, such mutants are true tolerance mutants, since a tolerance phenotype occurs when initial resistance is unaffected. Furthermore, they are also predicted to be true tolerance mutants by our correlation, since our curve is close to the origin of the plot (x = y = 0), and residuals at x = 0 (i.e., no initial resistance phenotype) equal the measured tolerance effect size y (see Figures 5B, 7A). *Kdm3* mutants show reduced initial resistance and reduced tolerance (counter to the prediction from the inverse correlation), suggesting that *Kdm3* might be involved in both setting initial neuronal activity and in the mechanism of increasing neuronal activity after the initial sedating ethanol exposure.

Conversely, the *HDM* mutant *No66* develops more tolerance than expected in 3 of 6 experiments. Interestingly, long exposures at low ethanol concentrations exacerbated tolerance strongly, while short, high-concentration exposures caused small effects on tolerance. This suggests that some mutant phenotypes are dependent on the kinetics of the exposure, but we did not observe this kinetic-dependent tolerance phenotype for other *HDM* mutants like *Kdm3.*^22^

Mutants with residuals far from zero provide potential insights into the mechanisms of alcohol tolerance. The residuals plotted as a histogram (Figure 7B) show a cluster of 6 data points with residuals of –3 or lower, suggesting that these mutants’ phenotypes reflect true tolerance mutants that develop little tolerance. These mutants include the genes encoding Gprk2, a G protein-coupled kinase,^21, 24^ CASK, a member of the MAGUK family of scaffolding proteins,^16^ the NMDA receptor, which interacts with CASK^25–27^, and Arf6, a small GTPase involved in membrane trafficking and actin remodeling and its activator Efa6.^28–30^ Mutants for Arf6 and Efa6 are highly sensitive to ethanol-induced sedation but induce very little tolerance response.^30^ We have placed Arf6 downstream of the insulin receptor (InR) and upstream of S6 kinase (S6k) in a pathway mediating initial resistance to ethanol. Overexpression of Arf6, as well as constitutive-active InR and S6k, causes increased initial resistance.^31^ Similarly, the loss of ArfGAP3, an inactivator of Arf6, also causes increased initial resistance. All of these manipulations lead to decreased tolerance (in percent, relative to the first exposure), but not more so than expected based on their initial resistance phenotypes. In other words, these manipulations cause phenotypes consistent with our described inverse correlation. This suggests that the activity of this pathway is necessary for the development of tolerance, while gains of function in this pathway are not sufficient to cause excessive tolerance. Similarly, mutants for RhoGAP18B and Rsu1, both negative regulators of Rho GTPases, affect initial resistance but not tolerance.^32, 33^ Mutations upstream of RhoGAP18B and Rsu1 in *mys*, encoding the cell adhesion molecule ß- integrin, are also initially sensitive but develop tolerance as expected (by the inverse correlation).^14^ This suggests that while both the Rho-family and Arf6 GTPase pathways are involved in setting initial resistance to ethanol exposure, only the Arf6 pathway is critical for ethanol-induced tolerance.

Although fewer data points have highly positive residuals, indicating the development of more tolerance than controls, mutant *2-10*^11^ and *No66*^22^ have residuals greater than 2. For *No66*, effects are dose-dependent, where a low ethanol concentration suggests that *No66* is a tolerance mutant while a high concentration does not. The relative lack of mutants which confer greater tolerance suggests that the biological mechanism of tolerance is more easily disrupted than potentiated. Additionally, these data suggest that LOF mutants may be more informative for studying ethanol tolerance since we did not see any GOF outliers that appeared to be true tolerance mutants.

A previous study has assessed the sedation and tolerance phenotypes of *Gprk2* mutants.^21^ These mutants were considerably more resistant and developed much less tolerance than control flies, consistent with the trend of our inverse correlation. Based on their findings, the authors of that study also wondered whether the *Gprk2* tolerance phenotype was a result of the initial resistance phenotype. To investigate this, they exposed *Gprk2* mutants and control flies to the same depth of sedation by exposing them to the ST-90, the time point at which 90% of the flies in each vial were sedated. After exposing flies to the ST-90, which took longer for *Gprk2* mutants, they still developed significantly less tolerance than controls and are, therefore, true tolerance mutants. This is also suggested by their residual of –4.85 from our correlation with the initial same-dose exposure data (Figure 7C). Exposure of both experimentals and controls to the ST-90 is conceptually similar to our *GNMT* experiments (Figure 6), where both groups were exposed to 1.5*ST-50, the result of both paradigms being that experimental and control flies are exposed to a consistent depth of sedation. These methods of determining *Gprk2*, but not *GNMT,* to be true tolerance mutants suggest that tolerance mutants may be most easily identified via a mechanism that exposes experimental and control flies to the same behavioral effect, such as the inebriometer.^34, 35^ In the inebriometer, flies are exposed to alcohol until they become sedated and elute out the bottom of the device, meaning that each fly is exposed to an equivalent dose of alcohol which confers a consistent behavioral effect. In the absence of an inebriometer, a similar exposure can be performed by exposing flies to 1.5*ST50, as we have done here, or to the ST-90, as performed by Kang and colleagues.^21^

Together, these findings suggest that initial resistance to alcohol sedation is a behavioral set point that impacts the subsequent development of tolerance. Accordingly, many mutants that are considered tolerance mutants may, in reality, be resistance mutants that appear to have tolerance phenotypes only because of their resistance phenotypes. Increasing the duration of the initial exposure increases the amount of tolerance developed.^1, 10^ Therefore, accurate assessment of potential tolerance mutants requires correcting for initial resistance phenotypes.

As mentioned above, a clear indication of a true tolerance mutant is that compared to the control, the mutant has a tolerance phenotype but no initial resistance phenotype (as is the case for mutants *2-10* and *5-21*, Figure 7A). However, when a mutant has both an initial resistance and tolerance phenotype, it is less clear whether it is a true tolerance mutant. Therefore, approaches to correcting for initial resistance would include testing potential tolerance mutants in an inebriometer or exposing experimental and control flies to the same depth of sedation with exposures of 1.5*ST-50 or to ST-90. However, as a preliminary step to performing these experiments, one can take advantage of the linear regression equation we have generated here (which is informed by 159 unique data points) to assess potential tolerance mutants. We determined our residuals using the linear regression equation (y = –0.56 x – 0.23). Because we used effect sizes for this correlation, they should be comparable from one experiment to the next regardless of the exact numerical naïve resistance and tolerance values.

Here, we provide a ‘how-to’ guide for using our linear regression equation to calculate residuals and assess potential tolerance mutants. This approach applies to experiments where experimental vs. control flies are exposed to a defined dose/duration of ethanol, with subsequent re-exposure to determine tolerance. First, determine the initial ST-50 and second (tolerance) ST-50 for mutant and control flies, as described previously.^12^ Then, calculate the effect sizes for resistance and tolerance (using, for example, this tool, where Group 1 is the control flies, and Group 2 is the experimental flies: https://www.psychometrica.de/effect_size.html. We recommend using #2, Hedge’s *g*, which controls for small sample sizes). Since the effect size is highly sensitive to the standard deviation, we recommend normalizing effect sizes if the standard deviation appears to be much lower or higher than is typical. Next, use our equation (y = –0.56 x –0.23), and plug in the initial resistance effect size for x. Then, subtract the result of this calculation from the effect size for tolerance measured experimentally. This will provide the residual for the mutant assayed.

As an example, we calculated the initial resistance and tolerance effect sizes for *GNMT*, *Arf6*, and *Gprk2* and plugged these into the equation. After subtracting the calculated values from their tolerance effect sizes, we found residuals of 0.18, –3.91, and –4.85, respectively (Figure 7C-D). These residuals support the conclusions that *GNMT* is not a tolerance mutant, while *Arf6* and *Gprk2* are both tolerance mutants, as determined by our findings and as reported previously.^21^ We suggest that utilizing our equation may be a useful starting point for determining promising candidate tolerance mutants to follow up on. Any mutants with absolute residuals from this equation around 2 or higher are most likely strong, true tolerance mutants, and any with absolute residuals between 1.3 and 2 may be worth investigating more closely. Should these candidates show a naïve resistance phenotype, this might include adjusting the first exposure to the same effective depth of sedation in subsequent experiments, as we have done here with genotypes exposed to 1.5*ST-50, or as done by Kang et al.,^21^ exposing to ST-90. Our data are thus widely applicable for investigating the mechanisms of alcohol-induced tolerance, an important but understudied endophenotype of AUD.^3^

## Abbreviations

AUD: Alcohol use disorder
CNS: Central nervous system
GOF: gain-of-function
HDM: Histone demethylase
LOF: loss-of-function
LORR: Loss of righting reflex
ST-50: Time to 50% sedation

## Acknowledgments

We thank members of the Rothenfluh and Rodan labs for continued discussion. Stocks obtained from the Bloomington Drosophila Stock Center (NIH P40OD018537) were used in this study. This work was supported by the NIH: grants F31AA030209 to M.M.C, K01AA029200 to C.B.M., R01DK110358 to A.R.R., and R01AA026818 & R01AA019526-S1 to A.R.

